# Platinum-Induced Mitochondrial OXPHOS Contributes to Cancer Stem Cell Enrichment in Ovarian Cancer

**DOI:** 10.1101/2022.02.01.478738

**Authors:** Shruthi Sriramkumar, Riddhi Sood, Thomas D. Huntington, Ahmed H. Ghobashi, Weini Wang, Kenneth P. Nephew, Heather M. O’Hagan

## Abstract

**Background:** Platinum based agents – cisplatin and carboplatin in combination with taxanes are used for the treatment of ovarian cancer (OC) patients. However, the majority of OC patients develop recurrent, platinum resistant disease that is uniformly fatal. Acute platinum treatment enriches for chemoresistant aldehyde dehydrogenase (ALDH) + ovarian cancer stem cells (OCSCs), which contribute to tumor recurrence and disease relapse. Acquired platinum resistance includes metabolic reprograming and switching to oxidative phosphorylation (OXPHOS). Chemosensitive cells rely on glycolysis while chemoresistant cells have the ability to switch between glycolysis and OXPHOS, depending on which pathway drives a selective advantage for growth and chemoresistance. High expression of genes involved in OXPHOS and high production of mitochondrial ROS are characteristics of OCSCs, suggesting that OCSCs favor OXPHOS over glycolysis. Based on connections between OCSCs, chemoresistance and OXPHOS, we hypothesize that platinum treatment induces changes in metabolism that contribute to platinum-induced enrichment of OCSCs.

**Methods:** The effect of cisplatin on mitochondrial activity was assessed by JC1 staining and expression of OXPHOS genes by quantitative RTPCR. Cisplatin-induced changes in Sirtuin 1 (SIRT1) levels and activity were assessed by Western blot. Small molecule inhibitors of mitochondrial complex I and SIRT1 were used to determine if their enzymatic activity contributes to the platinum-induced enrichment of OCSCs. The percentage of ALDH+ OCSCs in OC cells and tumor tissue from xenograft models across different treatment conditions was analyzed using ALDEFLUOR assay and flow cytometry.

**Results:** We demonstrate that acute platinum treatment increases mitochondrial activity. Combined treatment of platinum agents and OXPHOS inhibitors blocks the platinum-induced enrichment of ALDH+ OCSCs *in vitro* and *in vivo*. Furthermore, platinum treatment increases SIRT1 levels and subsequent deacetylase activity, which likely contributes to the increase in platinum-induced mitochondrial activity.

**Conclusions:** These findings on metabolic pathways altered by platinum-based chemotherapy have uncovered key targets that can be exploited therapeutically to block the platinum-induced enrichment of OCSCs, ultimately improving the survival of OC patients.

## Background

Ovarian cancer (OC) is the 5^th^ leading cause of cancer related deaths among women. The five-year survival rate of OC patients has remained low for decades at 46% (1). Surgical debulking followed by platinum and taxane-based chemotherapy are standard treatment modalities for OC patients (2). The most common histologic sub-type of serous epithelial OC – high grade serous epithelial OC (HGSOC) initially responds well to platinum-based chemotherapy (3). However, eventually patients with HGSOC relapse and develop a chemoresistant disease. Development of platinum resistance is the major obstacle in improving the survival of OC patients.

In the majority of solid tumors including OC, it has been firmly established that cancer stem cells (CSCs) are responsible for the development of chemoresistance and disease recurrence (4–6). Aldehyde dehydrogenase (ALDH) is the most commonly used robust marker of CSCs (6), and ALDH+ OCSCs have high tumor initiation capacity, enhanced ability to grow as spheroids and express high levels of stemness genes like *BMI1, OCT4 and NOTCH3* (7, 8). Furthermore, high expression and activity of the ALDH1A isoform strongly correlates with platinum resistant OC cells (8–10) and negatively correlates with survival of OC patients (11, 12). Our group as well as others have demonstrated that acute treatment of OC cells with platinum-based agents results in enrichment of ALDH+ OCSCs (7, 13, 14). Collectively, the evidence suggests that platinum-induced enrichment of ALDH+ OCSCs contributes to OC recurrence and relapse.

OCSCs are resistant to glucose deprivation and primarily rely on oxidative phosphorylation (OXPHOS) for energy requirements, which may also contribute to chemotherapy resistance (15–17). Chemosensitive OC cells favor glycolysis while chemoresistant OC cells are able to switch between glycolysis and OXPHOS depending on which pathway provides selective advantage for growth and platinum resistance (16). OCSCs exhibited high mitochondrial reactive oxygen species (ROS) production and high expression of enzymes involved in OXPHOS, implying that OCSCs preferentially utilize mitochondrial OXPHOS (15). Lung and pancreatic CSCs also displayed higher membrane potential and lower glucose consumption rates compared to non-CSCs (18, 19). Thus, there is strong evidence supporting altered metabolism as an important factor driving platinum resistance.

ALDH is a nicotinamide adenine dinucleotide (NAD+) dependent enzyme (20, 21). We and others have demonstrated that acute platinum treatment results in an increase in expression of nictotinamide phosphoribosyltransferase (NAMPT) – a rate limiting enzyme in NAD+ biosynthesis salvage pathway. The subsequent increase in cellular NAD+ levels promotes ALDH+ OCSC enrichment and contributes to platinum resistance (13, 14). However, it is well-established that platinum resistance is a multifactorial phenomenon (22) and it likely that additional metabolic pathways contribute to the enrichment of ALDH+ OCSCs and chemoresistance.

In the present study, we examined the effect of acute platinum treatment of HGSOC cells on mitochondrial activity. We observed that mitochondrial membrane potential (ΔΨ_M_- a surrogate for OXPHOS) and expression of genes involved in mitochondrial OXPHOS increased 16 hours after platinum treatment. Concomitantly, expression of genes in the glycolysis pathway was decreased by acute platinum treatment. Treatment of OC cells with OXPHOS inhibitors blocked the acute platinum-induced enrichment of ALDH+ OCSCs, and increased deacetylase activity of sirtuin 1 (SIRT1) was required for platinum-induced enrichment of ALDH+ OCSCs. In addition, we observed that SIRT1-regulated expression of mitochondrial transcription factor A (TFAM) likely contributed to the platinum mediated increase in mitochondrial activity. This first report demonstrating that OXPHOS inhibitors block platinum-induced enrichment of OCSCs supports further investigation into the role of mitochondrial OXPHOS in ovarian tumor recurrence.

## Methods

### Cell Culture

HGSOC cell lines used in the study were maintained at 37°C and 5% CO_2_ as described previously (23). OVCAR5 and OVSAHO cell lines were authenticated by ATCC in 2018. Briefly, OVCAR5 and OVSAHO cells were cultured in DMEM 1X (Corning, #MT10013CV) and RPMI 1640 (Corning, #MT10040CV), respectively, containing 10% FBS (Gibco, #16000044) without antibiotics. All the cell lines used in the study were passaged less than 15 times. A 1.67 mM stock solution of cisplatin (Millipore Sigma, #232120) was made using 154 mM NaCl (Macron Fine chemicals, #7581-12) in water. Stock solutions of Rotenone (Sigma, #R8875-1G; 100 mM), IACS – 010759-OXPHOS Inhibitor (MedChemExpress, #HY-112037; 10 mM), oligomycin (SelleckChem, #S1478; 10mM), and SIRT1 inhibitor Ex- 527 (Sigma, E7034; 10 mM) were made in DMSO. For all the experiments using these inhibitors, an equivalent amount of DMSO or inhibitors were added along with cisplatin and cells were incubated for 16 hours at 37°C and 5% CO_2_. All the treatment doses are specified in the figure legends.

### JC-1 Staining

OVCAR5 (1 × 10^6^) and OVSAHO cells (1.5 × 10^6^) were cultured in 100 mM plates for 24 hours and treated with respective IC_50_ doses of cisplatin (OVCAR5 and OVSAHO 12 μM and 4 μM, respectively (24)) for 16 hours. JC-1 staining was performed as per the manufacturer’s protocol (Thermo Fisher, #T3168). Briefly, cells were collected and washed with PBS and then resuspended in 1 ml PBS. Stock solutions of JC-1 were made in DMSO at a concentration of 5 mg/ml. Then JC-1 stain was added to cells at a final concentration of 2 μg/ml and cells were incubated at 37°C, 5% CO_2_ for 30 minutes. Following incubation, the cells were filtered through 30 μm filters and analyzed by flow cytometry.

### RNA isolation and quantitative reverse transcription PCR (qRT-PCR)

Isolation of total RNA from cell pellets was performed using RNAeasy mini kit (Qiagen, #74104) as per the manufacturer’s protocol. Maxima first strand cDNA synthesis kit (Thermo Fisher, #K1642) for quantitative reverse transcription PCR was used to synthesize cDNA. FastStart Essential DNA green master (Roche, #06402712001) was used to perform qRT-PCR. Primers of all candidate genes are in Supplemental Table 1. Expression of all the candidate genes were normalized to the housekeeping gene *Actin B*.

### ALDEFLUOR assay

OVCAR5 (1.0 ×10^5^) and OVSAHO (1.5 ×10^6^) were cultured in 100 mm dishes. Approximately 24 hours after plating, cells were treated with cisplatin alone or in combination with DMSO or appropriate inhibitors and incubated at 37°C, 5% CO_2_ for 16 hours. After the incubation time ALDEFLUOR assay (Stem Cell Technologies, #01700) was performed as described previously (14).

### Flow Cytometry

LSR II Flow cytometer (BD Biosciences) was used for the analysis of both the JC-1 and ALDEFLUOR assay. JC1 J-monomers and J-aggregates was measured using 488nm excitation and signal was detected using AF488 (green) and PE-A (red), respectively. For all the ALDEFLUOR assays, 488 nm excitation was used and the signal was detected using 530/30 filter to measure ALDH activity. To determine the percentage of ALDH positive cells in different conditions, the respective DEAB negative control was used. All the data analysis was done using FlowJo software (Becton, Dickinson & Company).

### Generation of stable knockdown lines

For SIRT1 knockdown (Sigma, NM_012238, TRCN0000018981, TRCN0000018983) and EV TRC1, the lentiviral shRNA knockdown protocol from the RNAi consortium of the Broad Institute was used as described previously (23).

### Mitochondrial DNA (mtDNA) content analysis

Mitochondrial DNA content was quantified relative to nuclear DNA (nDNA) (β_2_ – microglobulin) using qPCR and Delta Ct method as described previously (25–27). See Supplementary Table 1 for primers.

### Xenograft studies

All the animal studies were performed in accordance with the Association for Assessment and Accreditation of Laboratory Animal Care International and approved by Indiana University Bloomington Institutional Animal Care and Use Committee as described previously (14). 2 × 10^6^ OVCAR3 cells were injected subcutaneously in the flanks of NSG mice (IUSCCC *In Vivo* Therapeutics Core). Once the established tumors reached >100mm^3^, they were randomized into 3 different treatment groups and treated with vehicle alone or combination of carboplatin and vehicle or IACS-010759 (OXPHOS inhibitor). Carboplatin was administered once a week intraperitoneally at 50 mg/kg for 3 weeks. IACS-010759 was prepared in 10% DMSO and 90% corn oil. IACS-010759 was administered orally at 7.5 mg/kg five days a week for 3 weeks. At the end of the treatment schedule, mice were sacrificed and tumors were collected, disassociated and used for ALDEFLUOR assay as we have done previously (7, 14). Tumor dissociation kit and gentle MACS dissociator (Miltenyi Biotech) was used to dissociate the tumors into single cells as per manufacturer’s protocol.

### Correlation analysis using cBioportal

Correlation analysis of expression of *SIRT1* and *TFAM* was performed using the cBioportal (28, 29) and RNA-seq data of ovarian cancer samples from The Cancer Genome Atlas (30).

### Antibodies

For western blot, the following antibodies were used: anti-pATMS1981 (Cell Signaling Technology (CST), #13050, 1:1000), anti-AcH4K16 (CST, #13534, 1:1000), anti-total H2A (CST, #12349, 1:1000), anti-SIRT1 (Santa Cruz, sc-74465, 1:1000) and anti-vimentin (CST, #5741).

### Statistics

The percentage of ALDH+ cells across different conditions after ALDEFLUOR assay was evaluated by one-way ANOVA with multiple comparisons using Graphpad prism and data is presented as ±SEM. Data of JC-1 staining, RT-qPCR and densitometry were evaluated by Student *t* test in Graphpad prism and excel.

## Results

### Acute platinum treatment increases mitochondrial OXPHOS activity

Development of platinum resistance in OC has been linked to increased reliance of cancer cells on OXPHOS (31). Therefore, first we determined if mitochondrial membrane potential (ΔΨ_M_- a surrogate for OXPHOS)(32) increased after treatment of OC cells with cisplatin. To measure ΔΨ_M_, a lipophilic, cationic dye called JC-1 was used. JC-1 enters the mitochondria of healthy cells with normal ΔΨ_M_ and spontaneously aggregates as red fluorescent J-aggregates; in contrast, in cells with disrupted ΔΨ_M_, JC-1 will not form aggregates and retains its green fluorescence (32). Treatment of HGSOC cells – OVCAR5, OVSAHO with their respective IC50 doses of cisplatin for 16 hours followed by JC-1 staining resulted in a shift in the population of cells from low red fluorescence to high red fluorescence (Figure 1A,B; Supplementary Figure S1A), indicating an increase in mitochondrial J-aggregates after acute cisplatin treatment. A histogram of the red fluorescence for JC-1 staining in untreated and cisplatin treated OVCAR5 and OVSAHO cells confirmed that cisplatin treated cells shifted towards higher red fluorescence (Figure 1C, D). Furthermore, the ratio of red to green fluorescence significantly increased after acute cisplatin treatment of both OVCAR5 and OVSAHO cells (Figure 1E, F). Treatment of OVCAR5 cells with cisplatin for 16 hours had no effect on the mitochondrial DNA (mtDNA) content relative to nuclear DNA content (Supplementary Figure S1B), further supporting that the cisplatin-induced increase in OXPHOS was due to an increase in mitochondrial activity, not an increase in mitochondrial number.

**Figure 1.**
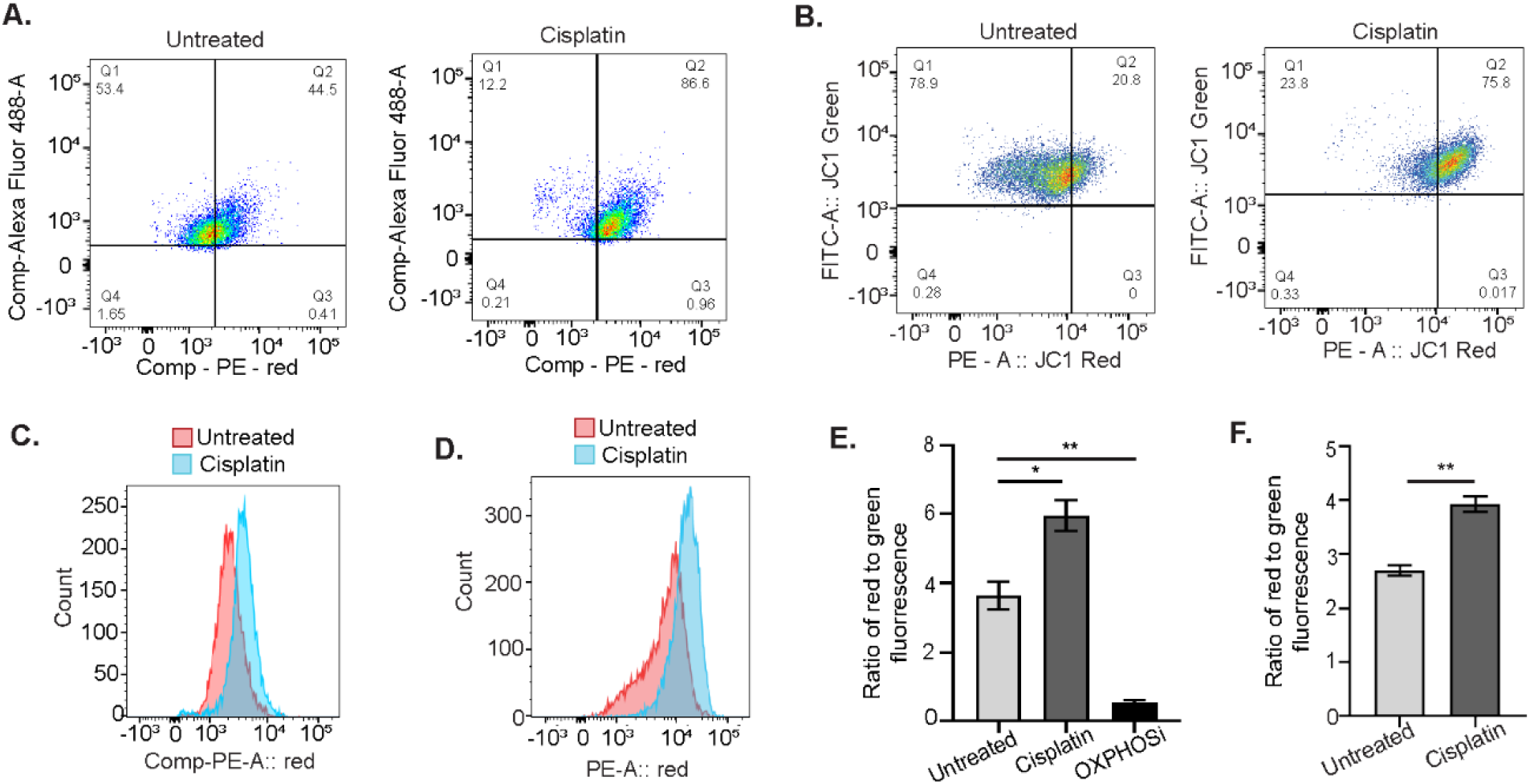
Mitochondrial OXPHOS activity increases in response to acute cisplatin treatment in OC cells. Scatter plots of OVCAR5 **(A)** and OVSAHO **(B)** cells untreated or treated with 12 μM and 4 μM cisplatin, respectively, for 16 hours followed by staining with JC-1 for 30 minutes and analysis by flow cytometry. Histogram of untreated and cisplatin treated OVCAR5 **(C)** and OVSAHO **(D)** cells after JC-1 staining as in A and B. Ratio of red to green fluorescence intensity in untreated and cisplatin treated OVCAR5 **(E)** and OVSAHO **(F)** cells after JC-1 staining as in A and B. The OXPHOS inhibitor carbonyl cyanide 3-chlorophenylhydrazone (OXPHOSi) was used as a control. Graphs display mean ratio of red to green fluorescence ± SEM (N=3). Student t-test was used to calculate statistical significance. For all untreated versus cisplatin treated, *P* values *< 0.05, ** < 0.005 and *** < 0.0005.

### Acute cisplatin treatment induces OXPHOS gene expression

To determine if acute cisplatin treatment increased expression of genes of mitochondrial OXPHOS complexes, we performed qRT-PCR of untreated and 16 hour cisplatin-treated cells. Increased expression of Complex I subunits - *NADH ubiquinone oxidoreductase subunits S6, A11 (NDUFS6, NDUFA11);* Complex III subunits – *ubiquinol-cytochrome C reductase subunit X, XI (UQCR10, UQCR11);* Cytochrome C oxidase complex subunits 5A, 6A *(COX5A, COX6A);* ATP synthase membrane subunit F *(ATP5MF)* and *translocase of the inner mitochondrial membrane (TIMM17A)* was observed in both OVCAR5 and OVSAHO cells after acute cisplatin treatment (Figure 2A, B). In addition, we examined genes associated with hexokinase, which is involved in the first step in glycolysis (33), as well as hypoxia-inducible factor 1 (HIF-1), known to inhibit mitochondrial biogenesis and respiration by negatively regulating the activity of transcription factor c-Myc (34, 35). Treatment of OVCAR5 and OVSAHO cells with cisplatin for 16 hours decreased expression of *hexokinase* isoforms *hexokinase 1* (*HK1)* and *hexokinase 2* (*HK2)* (Figure 2C, 2D) as well as *HIF1α* (Figure 2E) and increased *c-Myc* expression (Figure 2F). Altogether, our findings suggest that acute platinum treatment caused an increase in ΔΨ_M_ and OXPHOS.

**Figure 2.**
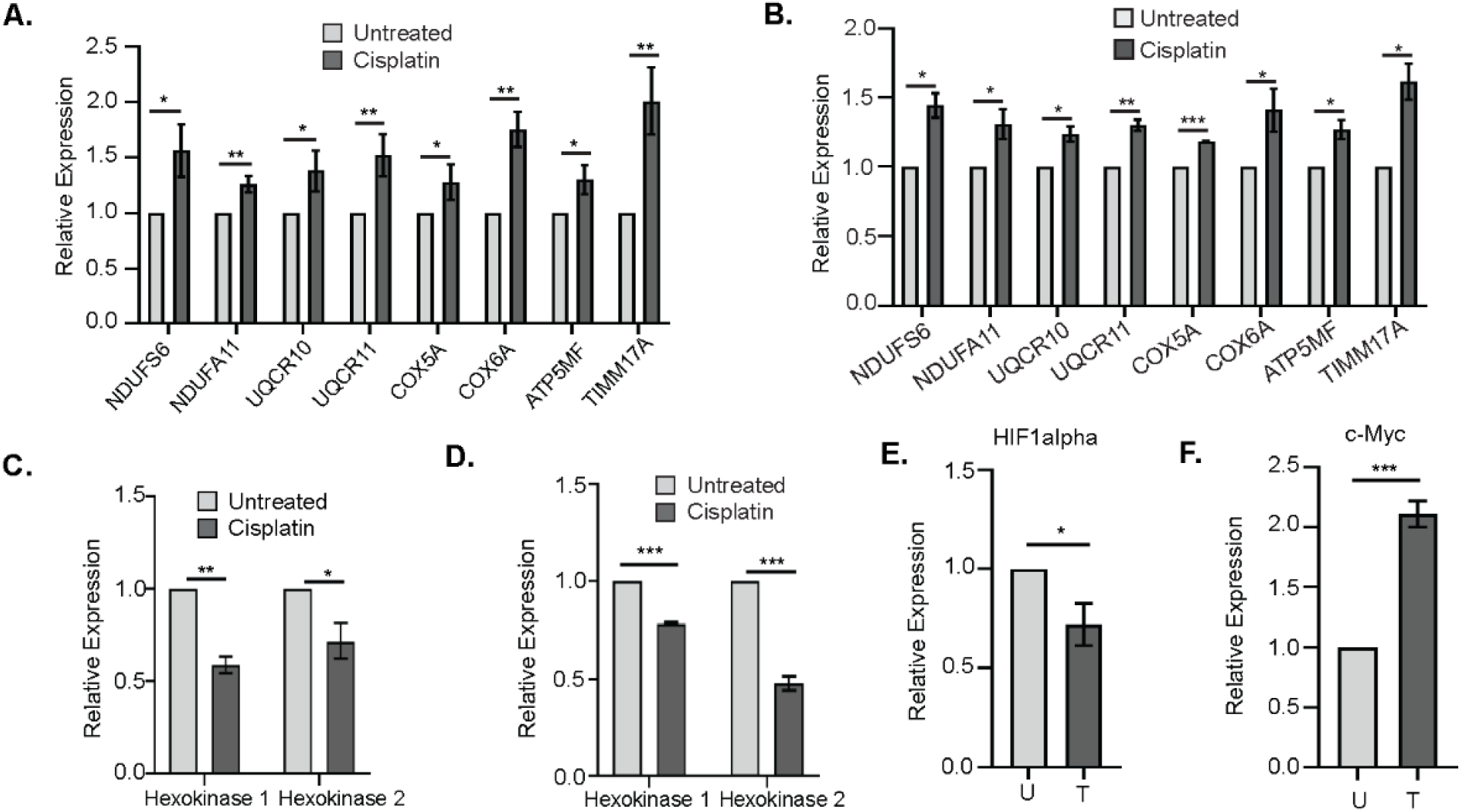
Expression of genes involved in mitochondrial OXPHOS increases after acute cisplatin treatment. Gene expression of the indicated genes by qPCR in OVCAR5 **(A)** and OVSAHO **(B)** cells untreated or treated with 12 μM or 4 μM cisplatin, respectively, for 16 hours. Expression of *hexokinase* isoforms in OVCAR5 **(C)** and OVSAHO **(D)** cells treated as in A and B. *HIF-1α* **(E)** and *c-Myc* **(F)** expression in OVCAR5 cells treated as in A. Graphs display mean fold change ±SEM relative to untreated. Expression of all the genes was normalized to the house keeping gene *Actin B*. For all untreated versus cisplatin treated, *P* values *< 0.05, ** < 0.005 and *** < 0.0005.

### OXPHOS is essential for the platinum-induced enrichment of ALDH+ cells

Acute platinum treatment has been previously demonstrated to enrich the ALDH+ OCSC population (7, 13, 14). To determine if the increase in OXPHOS in response to acute cisplatin treatment is essential for the platinum-induced enrichment of ALDH+ cells, we treated OVCAR5 and OVSAHO cells with cisplatin alone or in combination with 5μM Rotenone, a mitochondrial complex I inhibitor, or DMSO and performed the ALDEFLUOR assay. As expected, treatment of OVCAR5 and OVSAHO cells with cisplatin alone or in combination with DMSO increased the percentage of ALDH+ OCSCs (Figure 3A, B). Rotenone treatment blocked the platinum-induced increase in %ALDH+ OCSCs without changing the basal percentage of ALDH+ cells. IACS – 010759 is a novel small molecule OXPHOS inhibitor, which like rotenone also inhibits complex I of the mitochondrial respiratory electron transport chain (36). Treatment of OVCAR5 and OVSAHO cells with 1μM IACS-010759 in combination with cisplatin abrogated the platinum-induced enrichment of ALDH+ cells (Figure 3C-E; Supplementary Figure S2A, S2B). Oligomycin inhibits the ATP synthase complex in the mitochondrial respiratory electron transport chain (37), and combination treatment of 1 μM oligomycin plus cisplatin blocked the platinum-induced increase in ALDH+ cells (Figure 3F). Altogether, these data suggest that the increase in mitochondrial activity observed after acute cisplatin treatment is required for the platinum-induced enrichment of ALDH+ OCSCs

**Figure 3.**
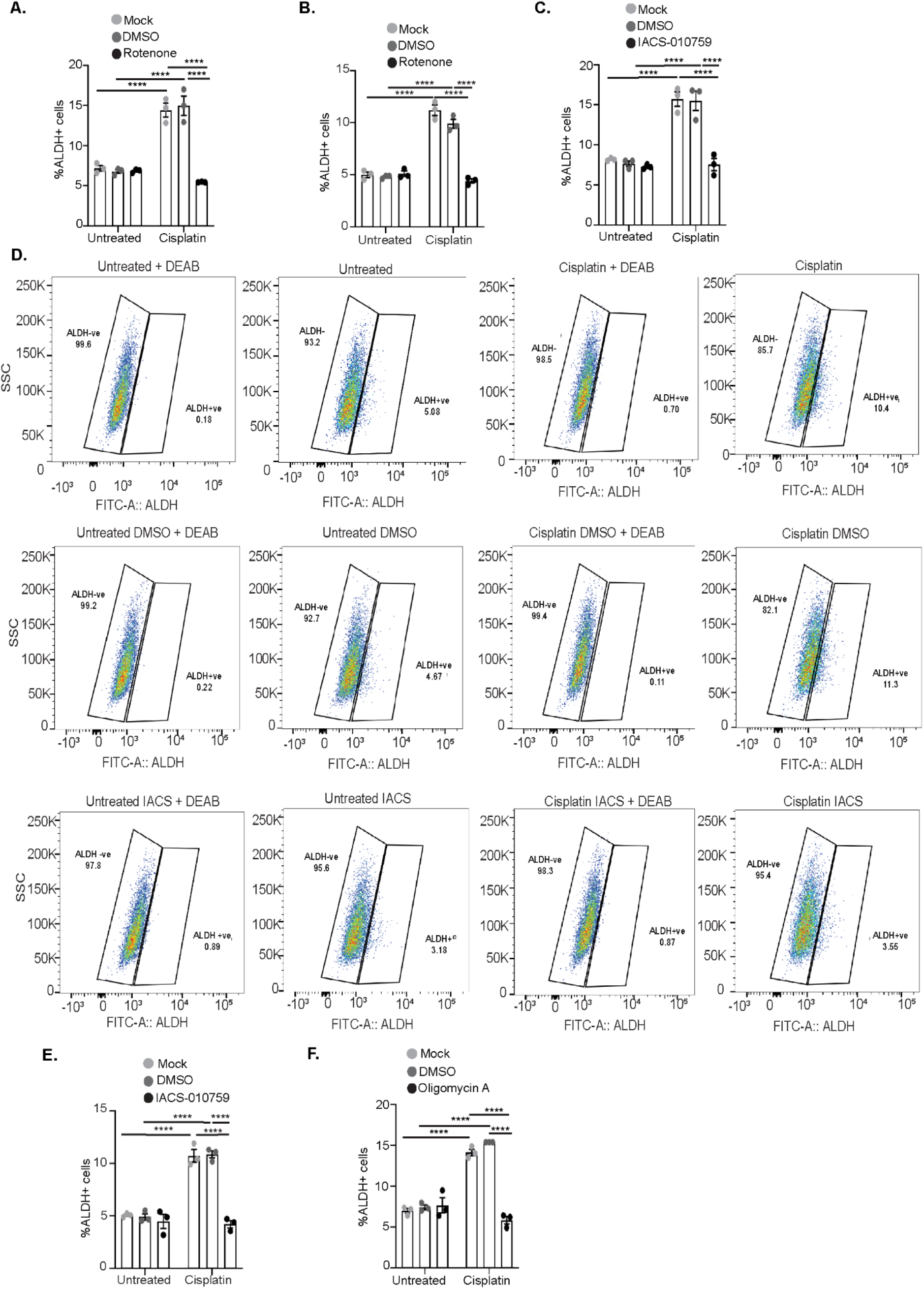
Mitochondrial OXPHOS inhibitors in combination with cisplatin block the platinum-induced increase in percent ALDH+ cells. Percent ALDH+ OVCAR5. **(A)** and OVSAHO cells **(B)** untreated or treated with 12 μM or 4 μM cisplatin, respectively, alone or in combination with DMSO or 5 μM Rotenone for 16 hours followed by ALDEFLUOR assay. Percentage ALDH+ OVCAR5 **(C)** and OVSAHO **(D-E)** cells treated with cisplatin alone or in combination with DMSO or 1 μM IACS-010759 for 16 hours followed by ALDEFLUOR assay. (D) shows gates used to determine ALDH+ cells using DEAB controls for one biological replicate of OVSAHO cells. **(F)** Percent ALDH+ OVCAR5 cells treated with cisplatin alone or in combination with DMSO or 1 μM oligomycin for 16 hours followed by ALDEFLUOR assay. Graphs display mean ± SEM percent ALDH+ cells in N = 3 biological replicates. For all untreated versus cisplatin treated, *P* values * < 0.05, ** < 0.005, ***< 0.0005, ****<0.0001.

### SIRT1 contributes to the platinum-induced enrichment of ALDH+ cells

SIRT1 deacetylates acetylated histone 4 lysine 16 (H4K16ac) as well as non-histone proteins and has been implicated in the regulation of metabolic processes (38, 39). Cellular NAD+ levels were reported to regulate the deacetylase activity of SIRT1 after acute platinum treatment (13, 14, 40), and SIRT1 expression and activity was also increased in platinum resistant OC cells (41, 42). However, the precise mechanism by which SIRT1 contributes to platinum resistance remains incompletely understood. Cisplatin treatment for 16 hours reduced the levels of H4K16ac relative to untreated OVCAR5 cells (Figure 4A). Phosphorylation of ATM (p-ATM) at serine 1981 was used as a control for platinum-induced DNA damage response and we observed an expected increase in ATM phosphorylation at S1981 after platinum treatment in OVCAR5 (Figure 4A). Treatment of OVCAR5 and OVSAHO cells with the IC50 dose of cisplatin for 16 hours also resulted in increased mRNA and protein levels of SIRT1 (Figure 4B-D). Furthermore, SIRT1 knockdown (KD) using shRNA rescued the platinum-induced decrease in H4K16Ac (Figure 4D), confirming that SIRT1 deacetylates H4K16 in response to acute cisplatin treatment. KD of SIRT1 using two different shRNAs in OVCAR5 cells followed by cisplatin treatment blocked the platinum-induced increase in ALDH+ cells and the baseline percentage of ALDH+ cells remained unchanged (Figure 4E). To further investigate if inhibiting SIRT1 would block platinum-induced enrichment of ALDH+ cells, OVCAR5 and OVSAHO cells were treated with cisplatin alone or cisplatin plus SIRT1 inhibitor Ex-527 (3 μM) or DMSO followed by ALDEFLUOR assay. In both OC cell lines, inhibiting SIRT1 completely abrogated the platinum-induced increase in ALDH+ cells (Figure 4F, G).

**Figure 4.**
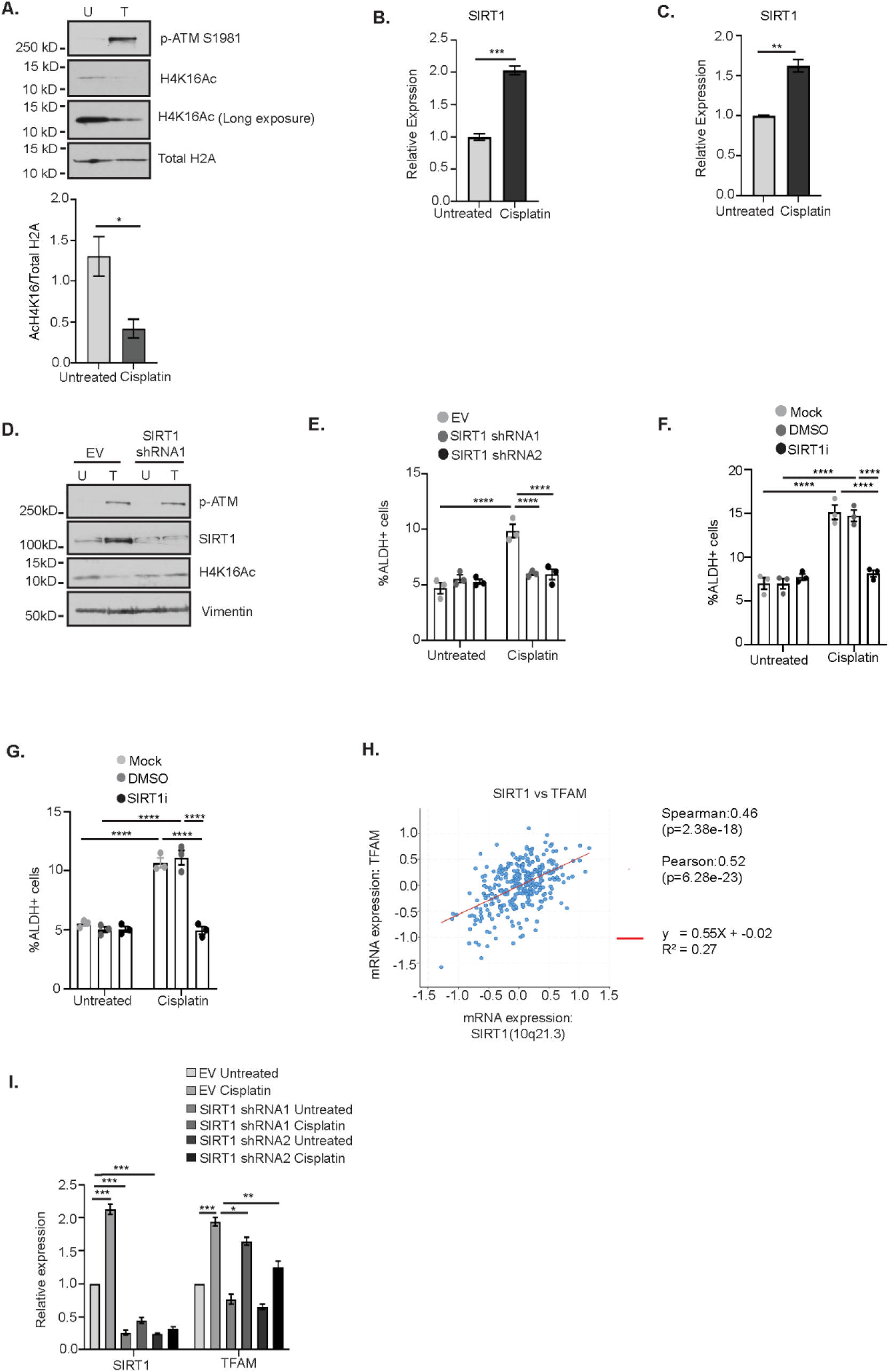
Platinum-induced increase in SIRT1 activity contributes to the enrichment of ALDH+ cells. **(A)** OVCAR5 cells were untreated (U) or treated with IC50 dose of cisplatin (12 μM) for 16 hours (T). Cell lysates were collected and analyzed by Western blot. Graph displays mean ±SEM densitometric analysis of N=3 biological replicates of H4K16ac relative to total H2A. Relative expression of SIRT1 in OVCAR5 **(B)** and OVSAHO **(C)** cells untreated or treated with 12 μM or 4 μM cisplatin, respectively, for 16 hours. **(D)** OVCAR5 cells infected with empty vector (EV) or SIRT1 viral shRNA followed by treatment as in C. Lysates were collected and analyzed by Western blot for the indicated proteins. **(E)** OVCAR5 cells infected EV or SIRT1 viral shRNA, untreated or treated as in C followed by ALDEFLUOR assay. Graph shows mean ±SEM percent ALDH+ cells in N=3 biological replicates. Percent ALDH+ OVCAR5 **(F)** and OVSAHO **(G)** cells treated with cisplatin alone or in combination with DMSO or 3 μM Ex-527 for 16 hours followed by ALDEFLUOR assay. Graph depicts mean ± SEM percent ALDH+ cells in N=3 biological replicates. **(H)** Correlation analysis of SIRT1 and TFAM expression using RNA-seq data from TCGA OC patient dataset. **(I)** Relative expression of SIRT1 and TFAM in OVCAR5 cells infected with EV or SIRT1 viral shRNA and treated as in C. Graph displays mean relative expression ±SEM to the untreated. Gene expression was normalized to the house keeping gene *Actin B*. For all comparisons, *P* – * < 0.05, **< 0.005, ***< 0.0005, ****<0.0001.

SIRT1 has been shown to promote OXPHOS and mitochondrial biogenesis in adipocytes and hepatocellular carcinoma cells by regulating the expression of mitochondrial transcription factor A (TFAM) (43–45). TFAM is important for proper transcription of mitochondrial and OXPHOS genes (46, 47). To test the hypothesis that SIRT1 promoted the platinum-induced increase in ALDH+ cells by promoting the increase in mitochondrial OXPHOS, we first analyzed the correlation between SIRT1 and TFAM expression in publicly available RNA-sequencing datasets from OC patients (30). SIRT1 expression was positively correlated with TFAM expression (Figure 4H). Next, we determined if the expression of TFAM is dependent on SIRT1 in response to acute cisplatin treatment in OC cells. SIRT1 KD using two different shRNAs resulted in a modest but significant reduction in the platinum-induced increase in TFAM expression (Figure 4I). Altogether, these data suggest that SIRT1 activity plays a key role in platinum-induced enrichment of ALDH+ cells in OC.

### OXPHOS inhibition blocks the platinum-induced enrichment of ALDH+ cells *in vivo*

We and others have demonstrated that acute platinum treatment results in enrichment of ALDH+ OCSCs *in vitro* and *in vivo* (7, 13, 48, 49). Furthermore, we demonstrated that treatment of OC cells with OXPHOS inhibitors in combination with cisplatin blocked the platinum-induced enrichment of ALDH+ cells *in vitro* without affecting the basal ALDH percentage (Figure 3A-F). Therefore, it was of interest to determine if OXPHOS inhibition blocks the platinum mediated increase in ALDH+ OCSCs *in vivo* in mouse xenografts. As shown in Figure 5A, 2 million OVCAR3 cells were injected subcutaneously in the flanks of NSG mice and after establishment of tumors, randomized mice received vehicle alone or a combination of carboplatin + vehicle or OXPHOS inhibitor IACS-010759. As expected, platinum treatment significantly reduced the tumor volume compared to the vehicle only group (Figure 5B). In the group treated with carboplatin + IACS-010759, tumor volume was less than the group treated with carboplatin alone at the end of the study (Figure 5B). We observed the expected increase in the percentage of ALDH+ cells in the carboplatin + vehicle compared to the vehicle only group, consistent with previous studies by us and others (7, 13, 48, 49). The combination treatment of IACS-010759 + carboplatin abrogated the platinum-induced enrichment of ALDH+ cells *in vivo* (Figure 5C). Altogether, we demonstrate that inhibiting OXPHOS with a complex I mitochondrial inhibitor blocks platinum-induced enrichment of ALDH+ cells *in vivo*.

**Figure 5:**
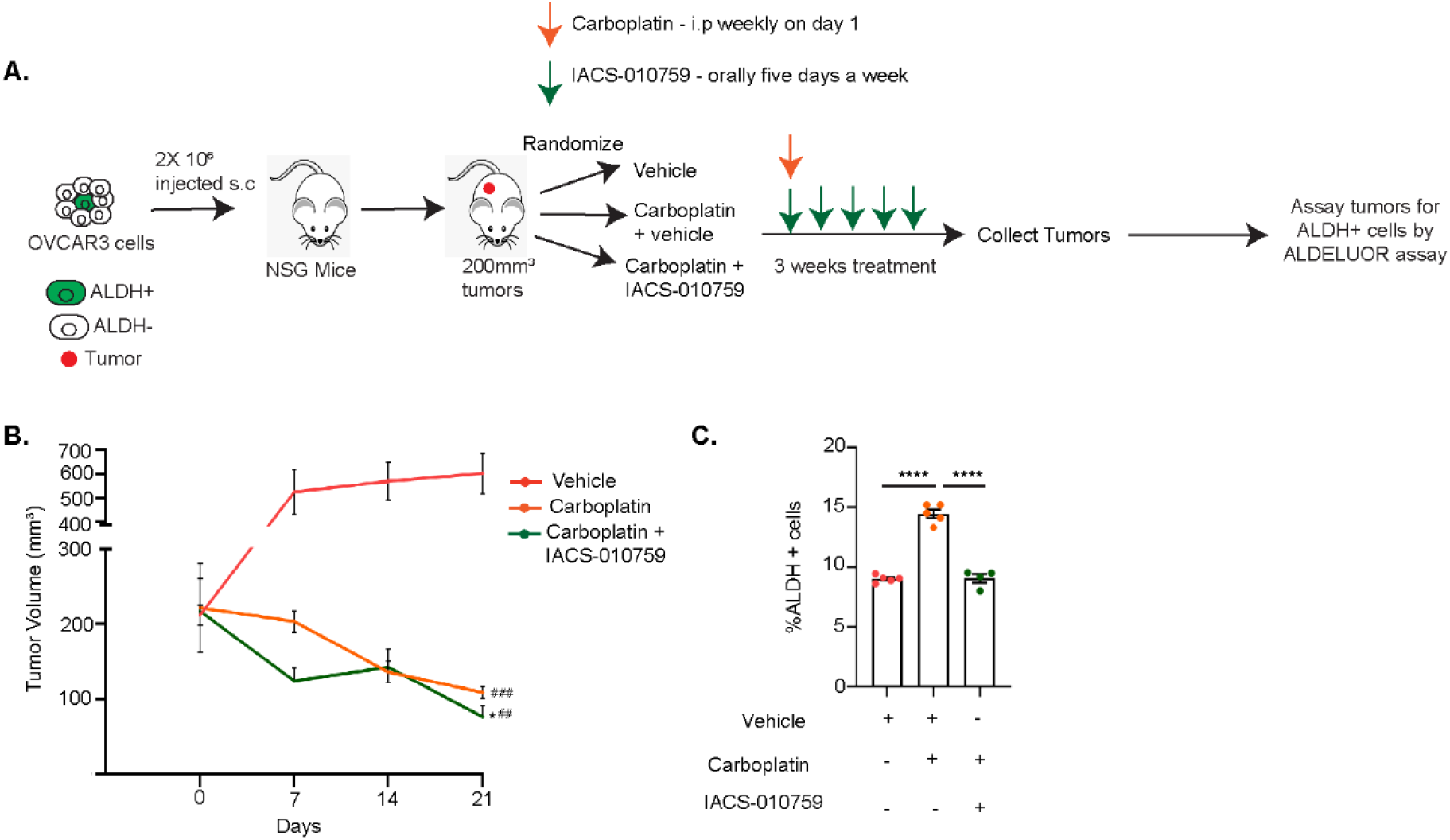
OXPHOS inhibition blocks the platinum-induced enrichment of ALDH+ cells *in vivo*. **(A)** 2 × 10^6^ OVCAR3 cells were injected s.c in 6-7 week old NSG mice. Once the established tumors were > 100mm^3^, mice were randomized into three groups and treated with vehicle alone or combination of vehicle + carboplatin or IACS-010759 for 3 weeks as indicated. At the end of the study, tumors were collected, dissociated into single cells and the ALDEFLUOR assay was performed. **(B)** Tumor volumes were measured using a digital caliper through 3 weeks of treatment. N=4-5 mice per group. \**P* relative to carboplatin + vehicle. #*P* relative to vehicle. **(C)** Percentage of ALDH+ cells in dissociated xenograft tumor samples using ALDEFLUOR assay. Graph indicates mean ±SEM ALDH percentage in the different treatment groups. N = 4-5 mice per group. For all treated with vehicle alone versus carboplatin + vehicle or IACS-010759, *P* values *< 0.05, **<0.005, ***<0.0005, ****<0.0001.

## Discussion

Metabolic reprogramming is a distinctive characteristic of chemoresistant OC cells and likely occurs as a consequence of metabolic adaptation to the tumor microenvironment (16). Metabolic reprogramming provides adenosine triphosphate (ATP) and precursors for macromolecular biosynthesis to meet the energy requirements for rapid proliferation and survival of tumor cells (50). In addition to promoting tumorigenesis, alterations in cellular metabolism foster resistance to chemotherapy in OC and other cancer types (51–53). The predominant mechanism of cytotoxicity of chemotherapeutic agents is inducing DNA damage and oxidative stress. Metabolic reprogramming enables tumor cells to adapt and manage pharmacological insults and oxidative stress (50). The precise mechanism of how metabolic reprogramming promotes chemoresistance in OC is not understood. Several studies have established that ALDH+ CSCs contribute to the development of chemoresistance in OC (7, 13, 14). Here, we demonstrate for the first time a role for OXPHOS and SIRT1 in the platinum-induced enrichment of ALDH+ OCSCs.

There has been a long-standing debate about whether CSCs use glycolysis or OXPHOS for survival (54). The metabolic pathway utilized by CSCs depends on which pathway provides a selective advantage in the tumor microenvironment. Here, we demonstrate that acute cisplatin treatment elevates mitochondrial OXPHOS concurrently with an increase in percentage of ALDH+ cells. Furthermore, OXPHOS inhibitors in combination with cisplatin block the enrichment of ALDH+ cells after acute cisplatin treatment, suggesting that the platinum mediated increase in mitochondrial activity is required for the enrichment of ALDH+ cells. These findings agree with Viale et al. that pancreatic CSCs depend on OXPHOS for survival (19, 55). It is likely that OXPHOS generates ROS to regulate metabolic plasticity of OCSCs, as recently suggested by Luo and Wicha (56).

Platinum resistant OC cells display elevated expression of the deacetylase SIRT1 (41, 42). We demonstrate that SIRT1 levels and activity increase after acute platinum treatment. NAD+ functions as a co-factor for several enzymes like ALDH and SIRT1 (21, 57), and the increase in SIRT1 activity and expression in response to acute cisplatin treatment may be due to platinum-induced increase in cellular NAD+ levels (13, 14). The dependency of the platinum mediated increase in ALDH+ cells on SIRT1 may be due to enhanced regulation of the platinum-induced metabolic switch to oxidative metabolism and improved antioxidant defenses by SIRT1, both of which could promote survival of CSCs (40, 58). SIRT1 supports mitochondrial function in muscle cells after exercise or starvation by regulating TFAM (59, 60), and we show that SIRT1 KD reduces the platinum-induced increase in *TFAM* expression. However, the precise mechanism of how SIRT1 contributes to the increase in ALDH+ cells and regulates TFAM in response to acute cisplatin treatment requires further investigation.

## Conclusions

The persistence of ALDH+ cells after chemotherapy is a major cause of disease recurrence and relapse in OC (4). We have demonstrated that acute cisplatin treatment of OC cells results in an increase in mitochondrial membrane potential. Our *in vitro* and *in vivo* data demonstrate that combination treatment with an OXPHOS inhibitor and platinum blocks the platinum-induced enrichment of ALDH+ cells. Our findings reinforce the need for additional preclinical and clinical investigations aimed at exploiting OXPHOS inhibitors to delay tumor recurrence and improve survival in OC.

## Supporting information

Supplementary Figures with legends

## List of Abbreviations

ΔΨ_M_: Mitochondrial membrane potential
ALDH: Aldehyde dehydrogenase
ATM: Ataxia telangiectasia mutated
ATP5MF: ATP synthase membrane subunit F
COX5A: Cytochrome C oxidase subunit 5A
COX6A: Cytochrome C oxidase subunit 6A
CSCs: Cancer stem cells
DMSO: Dimethyl sulfoxide
HGSOC: High grade serous ovarian cancer
HIF1α: Hypoxia-inducible factor 1
HK1: Hexokinase 1
HK2: Hexokinase 2
KD: Knockdown
mtDNA: Mitochondrial DNA
NAD+: Nicotinamide adenine dinucleotide
NAMPT: Nicotinamide phosphoribosyltransferase
NDUFA11: NADH ubiquinone oxidoreductase subunit 11
NDUFS6: NADH ubiquinone oxidoreductase subunit 6
OC: Ovarian Cancer
OCSCs: Ovarian Cancer Stem Cells
OXPHOS: Oxidative phosphorylation
qRT-PCR: quantitative reverse transcriptase polymerase chain reaction
ROS: Reactive oxygen species
shRNA: short hairpin RNA
SIRT1: Sirtuin 1
TFAM: Mitochondrial transcription factor A
TIMM17A: Translocase of the inner mitochondrial membrane
UQCR10: Ubiquinol- cytochrome C reductase subunit X
UQCR11: Ubiquinol-cytochrome C reductase subunit XI

## Declarations

### Ethics approval and consent to participate

The animal study was approved by Indiana University Bloomington Institutional Animal Care and Use Committee and was performed in adherence with the Association for Assessment and Accreditation of Laboratory Animal Care International.

## Consent for publication

Not applicable

## Availability of data and materials

All the data generated in the manuscript and materials used in the study are available upon reasonable request.

## Author Contributions

SS conceived, designed, analyzed all the experiments, wrote the original draft of the manuscript, edited and proofread the manuscript. RS, TDH and WW assisted in data acquisition. AHG performed some validation experiments and proofread the manuscript. KPN provided resources, funding acquisition, edited and proofread the manuscript. HMOH conceived, assisted with experimental design, supervised the study, and provided resources, funding acquisition, edited and proofread the manuscript. All authors have approved the submitted version of the manuscript.

## Acknowledgments

We thank Christiane Hassel and the Indiana University Flow Cytometry facility for their assistance.

