## Supplementary Figures with legends for "Platinum-Induced Mitochondrial OXPHOS Contributes to Cancer Stem Cell Enrichment in Ovarian Cancer"

### Supplementary Figure S1

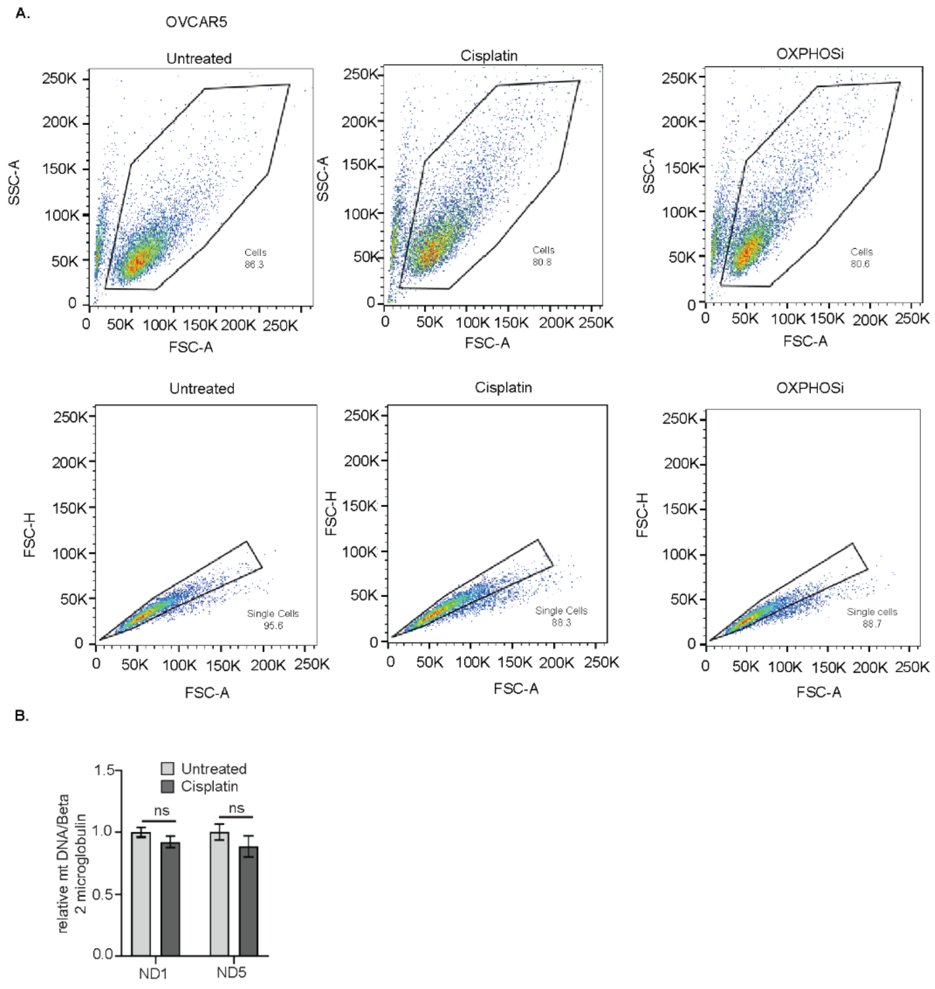


**Supplementary Figure S1 – mtDNA does not increase in response to acute cisplatin treatment. (A)** Plots of OVCAR5 cells untreated, OXPHOSi or 16 hr cisplatin after JC-1 staining. **(B)** OVCAR5 cells were treated with IC50 dose of cisplatin for 16 hours. DNA was isolated and qPCR was performed using two primer sets for different regions in the mitochondrial DNA relative to beta 2 microglobulin in the genomic DNA. Graph depicts mean ± SEM of mt DNA relative to nuclear DNA of N=3 biological replicates, *P* values *<0.05, **<0.005, ***<0.0005.

**Supplementary Figure S2**

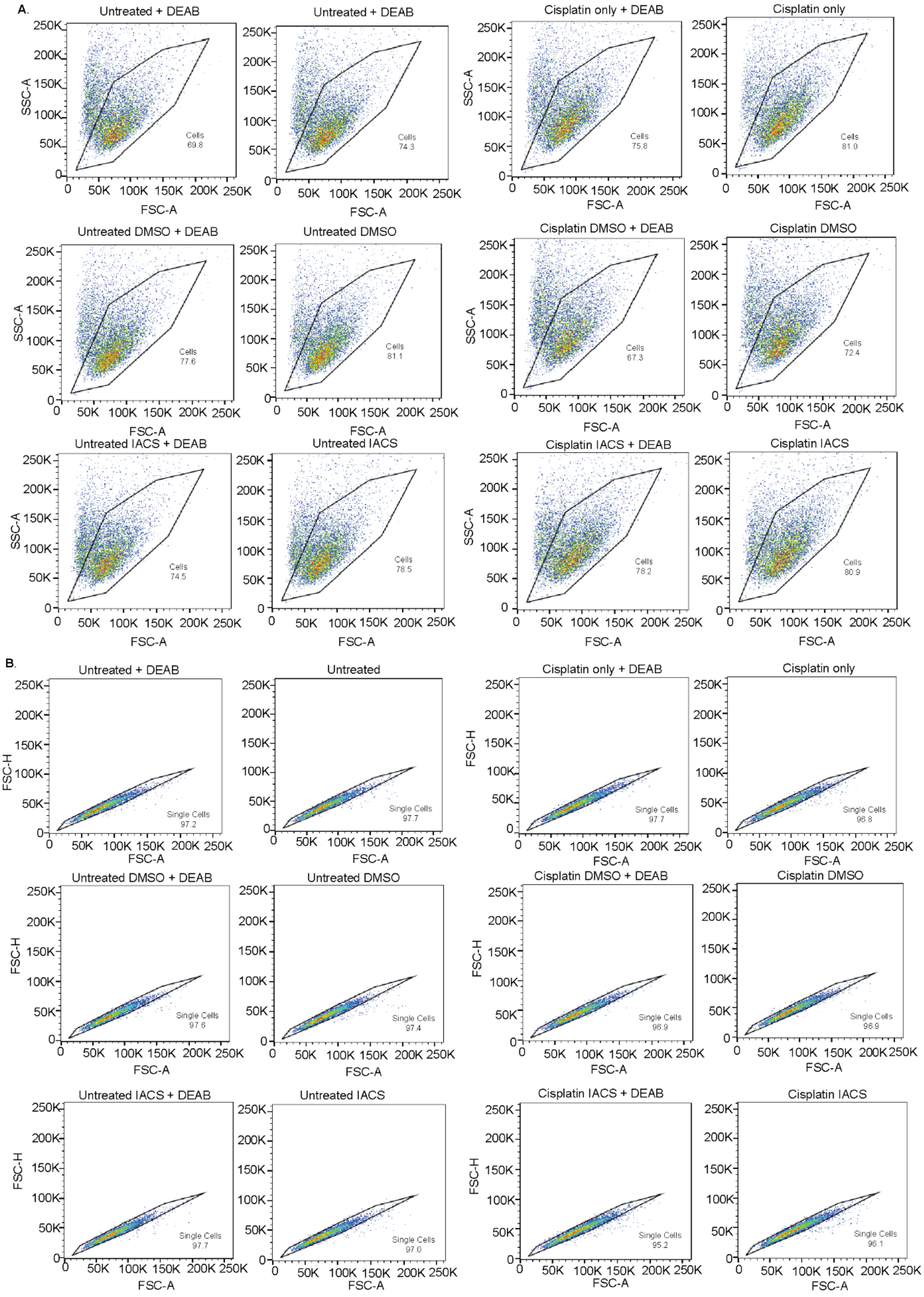
A.
Untreated + DEAB
Untreated + DEAB
Cisplatin only + DEAB
Cisplatin only
Untreated DMSO + DEAB
Untreated DMSO
Cisplatin DMSO + DEAB
Cisplatin DMSO
Untreated IACS + DEAB
Untreated IACS
Cisplatin IACS + DEAB
Cisplatin IACS
SSC-A
FSC-A
0
50K
100K
150K
200K
250K
Cells 69.8
Cells 74.3
Cells 75.8
Cells 81.0
Cells 77.6
Cells 81.1
Cells 67.3
Cells 72.4
Cells 74.5
Cells 78.5
Cells 78.2
Cells 80.9
B.
Untreated + DEAB
Untreated
Cisplatin only + DEAB
Cisplatin only
Untreated DMSO + DEAB
Untreated DMSO
Cisplatin DMSO + DEAB
Cisplatin DMSO
Untreated IACS + DEAB
Untreated IACS
Cisplatin IACS + DEAB
Cisplatin IACS
FSC-H
FSC-A
Single Cells 97.2
Single Cells 97.7
Single Cells 97.7
Single Cells 96.8
Single Cells 97.6
Single Cells 97.4
Single Cells 96.9
Single Cells 96.9
Single Cells 97.7
Single Cells 97.0
Single Cells 95.2
Single Cells 96.1

**Supplementary Figure S2 – Treatment of OC cells with mitochondrial complex I inhibitor in combination with cisplatin blocks platinum-induced enrichment of ALDH+ cells. (A-B)** Plots of ALDH percentage in OVSAHO cells treated with cisplatin alone or in combination with DMSO or 1µM IACS-010759 for 16 hours after ALDELUOR assay.
