## Supplementary Tables for "Platinum-Induced Mitochondrial OXPHOS Contributes to Cancer Stem Cell Enrichment in Ovarian Cancer"

**Table 1**

**qRT- Primer Sequences**

| Gene | Forward Primer | Reverse Primer |
|---|---|---|
| *NDUFS6* | TCTTGGCCACCCAAAAGTGT | CGGAAATGCTCACAGGATGC |
| *NDUFA11* | TGAACTACTTCCTCGGTGGC | CGCTATGCCAAAGTACACGC |
| *UQCR10* | TCGTGGGCGTCATGTTCTTC | GGGGGCCTCCAAGGAACTA |
| *UQCR11* | TCATCCTGAGGGTGCGACTC | GATCAGCCGCCAATCGGT |
| *COX5A* | TCATTGATGCTGCTTTGCGG | GTTCCTGGATGACATAGGGGT |
| *COX6A* | CATCTCCGCATCAGGACCAA | TCATCTTCGTAGCCAGTTGGA |
| *ATP5MF* | ACCAGGACTCCAAAATGGCG | CACTAGGACTGAAGTCCCGC |
| *TIMM17A* | GGTGGGGCCTTTACGATGG | GCCCTGGTTTTAATAGCTGTCA |
| *HIF1α* | TGAAGACATCGCGGGGAC | CTGGCTGCATCTCGAGACTTT |
| *c-Myc* | GGACCCGCTTCTCTGAAAGG | TAACGTTGAGGGGCATCGTC |
| *TFAM* | ACCAAAAAGACCTCGTTCAGC | TCAGAGTCAGACAGATTTTTCCAGT |
| *SIRT1* | TGACTGGACTCCAAGGCCACG | TCAGGTGGAGGTATTGTTTCCGGC |
| *Actin B* | GAAGCCGGCCTTGCACAT | AGCACAGAGCCTCGCCTTT |

**Primers for mtDNA content**

| Gene | Forward Primer | Reverse Primer |
|---|---|---|
| NADH dehydrogenase sub-unit 1 (ND1) | TTCTAATCGCAATGGCATTCCT | AAGGGTTGTAGTAGCCCGTAG |
| NADH dehydrogenase sub-unit 5 (ND5) | TTCATCCCTGTAGCATTGTTCG | GTTGGAATAGGTTGTTAGCGGTA |
| Beta 2 microglobulin | TGCTGTCTCCATGTTTGATGTATCT | TCTCTGCTCCCCACCTCTAAGT |
